# Development of a Nested-MultiLocus Sequence Typing approach for a highly sensitive and specific identification of *Xylella fastidiosa* subspecies directly from plant samples

**DOI:** 10.1101/2020.06.08.140194

**Authors:** Sophie Cesbron, Enora Dupas, Quentin Beaurepère, Martial Briand, Miguel Montes Borrego, Maria del Pilar Velasco Amo, Blanca B. Landa, Marie-Agnès Jacques

**Affiliations:** IRHS-WR1345, Université d’Angers, INRAE, Institut Agro, SFR 4207 QuaSaV, 49071, Beaucouzé, France; Institute for Sustainable Agriculture, Consejo Superior de Investigaciones Científicas (IAS-CSIC), 14004, Córdoba, Spain; French Agency for Food, Environmental and Occupational Health & Safety, Plant Health Laboratory, Angers, France

**Author notes:** **Correspondence:** Sophie Cesbron.

**Keywords:** *Xylella fastidiosa*, direct typing technique, MLST, nested PCR

## Abstract

Different sequence types (ST) of *Xylella fastidiosa* were already identified in France and Spain based on direct MultiLocus Sequence Typing (MLST) of plant DNA samples. However, direct typing of plant DNA is partly efficient. In order to improve the sensitivity of *X. fastidiosa* identification, we developed a direct nested-MLST assay on plant extracted DNA. This method was performed based on a largely used scheme targeting seven housekeeping gene (HKG) loci (*cysG, gltT, holC, leuA, malF, nuoL, petC*). Nested primers were designed from multi-sequence alignments of 38 genomes representing all subspecies and one genome of *Xylella taiwanensis*. Sequences obtained were long enough to be used for BLAST comparison in PubMLST database. No nonspecific amplification products were observed in these samples. Efficiency of the nested-MLST was tested on extracted DNA from 106 samples proven positive (Cq<35) or equivocal (35≤Cq≤40) using the Harper’s qPCR test. Samples analyzed included 49 plant species and two insect species (*Philaenus spumarius, Neophilaenus campestris*) that were collected in 2017 (106 plant samples in France), in 2018 (162 plant samples in France, 40 plant samples and 26 insect samples in Spain), and in 2019 (30 plant samples in Spain). With the conventional-MLST assay, no complete MLST profile was obtained for any of the samples from France and for most samples (59/66) from Spain. Conversely, with the nested approach, complete profiles were obtained for six French plant samples, 55 Spanish plant samples and nine Spanish insect samples. The threshold was improved by 100 to 1000 times compared to conventional PCR and was between 22 pg.mL^−1^ to 2.2 pg.mL^−1^ depending on the HKG. Using nested-MLST assay, plants that were not yet considered hosts tested positive and revealed novel alleles in France, whereas for Spanish samples it was possible to assign the subspecies or ST to samples considered as new hosts in Europe. Direct typing by nested-MLST from plant material has an increased sensitivity and may be useful for epidemiological purposes.

## Introduction

*Xylella fastidiosa* (*Xf*) is the causal agent of several devastating diseases of plants in the Americas and this pathogen was recently detected in Europe, where it causes a severe disease in olive tree in Italy and is present in several other regions. This species encompasses three well recognized subspecies, namely *fastidiosa, multiplex*, and *pauca* (Marcelletti and Scortichini, 2016; Denancé et al. 2019) but other subspecies are currently described (EFSA, 2018). The subspecies *fastidiosa* occurs in North and Central America and was recently detected in Spain (https://gd.eppo.int/taxon/XYLEFA/). It infects a wide range of trees, ornamentals, and other perennials and includes strains responsible for the well-known Pierce’s disease on grapevine (Janse & Obradovic 2010; EFSA 2018). The subspecies *multiplex* is present in North and South America and in Europe (https://gd.eppo.int/taxon/XYLEFA/) and is associated with scorches and dieback of a wide range of trees and ornamentals (EFSA 2018). The subspecies *pauca* is mostly found in South and Central America on *Citrus* spp. and *Coffea* spp. (Almeida et al. 2008), but has been recently detected also in olive trees in Spain (Landa, 2017), Brazil (Della Coletta-Filho et al. 2016), Argentina (Haelterman et al. 2015), and Italy (Saponari et al. 2013). Its host range includes also ornamentals and other trees (EFSA 2018). Altogether more than 560 plant species are hosts of *Xf* (EFSA 2018). This member of the *Xanthomonadaceae* family inhabits the xylem of its host plants (Wells et al. 1987) and is naturally transmitted by insects from plants to plants.

A range of detection tests has been proposed for *Xf* (EPPO 2019). Several immunological methods are available (EPPO 2019). However, such methods have high limits of detection (LoDs) that are close to 10^4^ to 10^5^ cells.mL^−1^ (EPPO 2019). End point and also quantitative PCR (qPCR) are nowadays widely used, with a better sensivity as the LoD is around 10^2^ cells.mL^−1^ for several qPCR tests (Ouyang et al. 2013, Harper et al. 2010, Bonants et al. 2019, Waliullah et al. 2019). The Harper’s qPCR test is often used in Europe for its high sensitivity, its target is located in the gene coding for the 16S rRNA-processing RimM protein. Several tests based on isothermal amplification have also been reported (Harper et al. 2010, Yaseen et al. 2015, Li et al 2016, Burbank & Ortega 2018, Waliullah et al. 2019). The Harper’s test has also been successfully transferred to be used in digital PCR (Dupas et al. 2019a). Some of these tests were designed to detect only one subspecies. This is the case of the nested PCR test proposed by Ciapina et al (2004) for detecting CVC strains (subspecies *pauca*) in sharpshooters and citrus plants and also of the qPCR test targeting oleander leaf scorch strains (that are included in the subspecies *fastidiosa*) (Guan et al. 2013). Other tests were designed to detect and discriminate two or more subspecies (Burbank & Ortega 2018; Dupas et al. 2019b).

Precise identification of *Xf* at an infraspecific level is essential for epidemiological and surveillance analyses, and to allow a proper description of the population structure and their dynamics. The widely used multilocus sequence typing (MLST) scheme designed for *Xf* (Scally et al. 2005, Yuan et al. 2010) is based on amplification by conventional PCR and sequencing of seven HKG fragments (loci), either from strains or from plant samples (Denancé et al., 2017). For each locus, the different sequence variants are considered as distinct alleles. The combination of allele numbers defines the sequence type (ST). The MLST-*Xf* data are stored in a public database (https://pubmlst.org/xfastidiosa/) that can be used to automatically identify and assign new allele variants, and provide tools to analyze the potential origin of the strains. The association of the different subspecies with their host plants is useful to better understand *Xf* epidemiology.

A reliable and enough informative typing method is particularly relevant in cases of new outbreaks or for the description of new host. Due to the large number of host plants to be analyzed, various types of inhibitors can interfere with reagents of PCR and low bacterial loads compromising PCR efficiency and hence typing. Improving DNA extraction methods can, at least partly, solve the problem of PCR inhibitors, and nested PCR appears a solution to allow the detection of low bacterial population sizes. A nested-MLST was already successfully developed to detect and type *Xf* in vectors (Cruaud et al. 2018). Primers were designed inside the gene fragments used in the conventional-MLST scheme and hence some informative sites are lost. MLST with nested PCRs has also been developed in medical field to enable the direct typing of samples infected by *Leptospira* or *Trichomonas*, for example (Weiss et al. 2016; Van der Veer et al. 2016).

The objective of this study was to develop a *Xf* detection assay based on the largely used MLST scheme (Yuan et al. 2010) that lowers the limit of detection (LoD) to enable at least the identification of *Xf* subspecies and, if possible, provide larger sets of typing data directly from plant samples. We used genomic sequences to improve each PCR efficiency and showed a drastic increase in the sensitivity as compared to that of the conventional-MLST approach.

## Materials and Methods

### Strains and media

A collection of target and non-target bacterial strains was used to test *in vitro* the specificity of the newly designed primers and the nested PCR assays. This set was made of five *X. fastidiosa* strains from different subspecies and 33 strains representing bacteria phylogenetically close to *Xf*, i.e. various *Xanthomonas*, as well as strains of other plant pathogenic bacteria and endosymbionts potentially inhabiting the same niches as *Xf* (Table 1), available at the French Collection of Plant-Associated Bacteria (CIRM-CFBP; https://www6.inra.fr/cirm_eng/CFBP-Plant-Associated-Bacteria). The *Xf* strains were grown on modified PWG media (agar 12 g.L^−1^; soytone 4 g.L^−1^; bacto tryptone 1 g.L^−1^; MgSO_4_.7H_2_O 0.4 g.L^−1^; K_2_HPO_4_ 1.2 g.L^−1^; KH_2_PO_4_ 1 g.L^−1^; hemin chloride (0.1% in NaOH 0.05 M) 10 ml.L-1; BSA (7.5%) 24 ml.L^−1^; L-glutamine 4 g.L^−1^) at 28°C for one week. *Agrobacterium* and *Rhizobium* were grown at 25°C for one to two days on MG medium (Mougel et al. 2001); TSA was used (tryptone soybroth 30 g.L^−1^; agar 15 g.L^−1^) for *Clavibacter*, *Ensifer*, *Stenotrophomonas*, *Xanthomonas* and *Xylophilus*; and King’s medium B (King et al. 1954) for *Dickeya*, *Erwinia*, *Pantoea* and *Pseudomonas*. For PCRs, bacterial suspensions were prepared from fresh cultures in sterile distilled water, adjusted at OD_600 nm_ = 0.1 and used as templates for amplification after boiling for 20 minutes, thermal shock on ice and centrifugation 10 000g, 10 min.

**Table 1:** List of target and non-target strains used to verify the specificity of nested-MLST primers

| CFBP code | Bacterial species | Host plant | Origin |
| --- | --- | --- | --- |
| 6448 | *Agrobacterium rubi* | *Rubus ursinus* var. *loganobaccus* | USA (1942) |
| 2413 | *Agrobacterium tumefaciens* | *Malus* sp. | NA (1935) |
| 5523 | *Agrobacterium vitis* | *Vitis vinifera* | Australia (1977) |
| 2404 | *Clavibacter insidiosus* | *Medicago sativa* | USA (1955) |
| 4999 | *Clavibacter michiganensis* | *Lycopersicon esculentum* | Hungary (1957) |
| 3418 | *Curtobacterium flaccumfaciens* pv. *flaccumfaciens* | *Phaseolus vulgaris* | Hungary (1957) |
| 1200 | *Dickeya dianthicola* | *Dianthus caryophyllus* | United Kingdom (1956) |
| 5561 | *Ensifer meliloti* | *Medicago sativa* | VA, USA (1984) |
| 1232 | *Erwinia amylovora* | *Pyrus communis* | United Kingdom (1959) |
| 3845 | *Pantoea agglomerans* | *Knee laceration* | Zimbabwe (1956) |
| 3167 | *Pantoea stewartii* pv. *stewartii* | *Zea mays* var. *rugosa* | USA (1970) |
| 3205 | *Pseudomonas amygdali* | *Prunus amygdalus* | Greece (1967) |
| 8305 | *Pseudomonas cerasi* | *Prunus cerasus* | Poland (2007) |
| 7019 | *Pseudomonas congelans* | na<sup>1</sup> | Germany (1994) |
| 1573 | *Pseudomonas syringae* pv. *persicae* | *Prunus persica* | France (1974) |
| 1392 | *Pseudomonas syringae* pv. *syringae* | *Syringa vulgaris* | United Kingdom (1950) |
| 7436 | *Rhizobium nepotum* | *Prunus ceresifera myrobolan* | Hungary (1989) |
| 13100 | *Stenotrophomas maltophilia* | *Phaseolus vulgaris* | Cameroon (2009) |
| 3371 | *Xanthomonas euvesicatoria* pv. *citrumelonis* | *Citrus* sp. | USA (1989) |
| 2528 | *Xanthomonas arboricola* pv. *juglandis* | *Juglans regia* | New Zealand (1956) |
| 2535 | *Xanthomonas arboricola* pv. *pruni* | *Prunus salicina* | New Zealand (1953) |
| 4924 | *Xanthomonas axonopodis* pv. *axonopodis* | *Axonopus scoparius* | Colombia (1949) |
| 5241 | *Xanthomonas campestris* pv. *campestris* | *Brassica oleracea* var. *gemmifera* | United Kingdom (1957) |
| 2901 | *Xanthomonas citri* pv. *aurantifolii* | *Citrus limon* | Argentina (1988) |
| 2525 | *Xanthomonas citri* pv. *citri* | *Citrus limon* | New Zealand (1956) |
| 7660 | *Xanthomonas citri* pv. *viticola* | *Vitis vinifera* | India (1969) |
| 2625 | *Xanthomonas gardneri* | *Medicago sativa* | Reunion Island (1986) |
| 4925 | *Xanthomonas hortorum* pv. *hederae* | *Hedera helix* | USA (1944) |
| 2533 | *Xanthomonas hortorum* pv. *pelargonii* | *Pelargonium peltatum* | New Zealand (1974) |
| 1156 | *Xanthomonas hyacinthi* | *Hyacinthus orientalis* | Netherlands (1958) |
| 2532 | *Xanthomonas oryzae* pv. *oryzae* | *Oryza sativa* | India (1965) |
| 2054 | *Xanthomonas translucens* | *Hordeum vulgare* | USA (1933) |
| 2543 | *Xanthomonas vasicola* pv. *holcicola* | *Sorghum vulgare* | New Zealand (1969) |
| 7970 | *Xylella fastidiosa* subsp. *fastidiosa* | *Vitis vinifera* | USA (1987) |
| 8416 | *Xylella fastidiosa* subsp. *multiplex* | *Polygala myrtifolia* | France (2015) |
| 8084 | *Xylella fastidiosa* subsp. *morus* | *Morus alba* | USA (na<sup>1</sup>) |
| 8070 | *Xylella fastidiosa* subsp. *multiplex* | *Prunus* spp. | USA (2004) |
| 8402 | *Xylella fastidiosa* subsp. *pauca* | *Olea europea* | Italy (2014) |

<sup>1</sup>: not available

### DNA extraction

Genomic DNA from *Xf* strain CFBP 8070 was extracted with the Wizard genomic DNA Purification Kit (Promega, France) and used to prepare a 10-fold serial dilutions from 220 ng.mL^−1^ (corresponding to 0.8×10^8^ copies.mL^−1^ of genomic DNA) to 22 fg.mL^−1^ (8 copies.mL^−1^) to evaluate the LoD of the nested-MLST. Copies number were calculated using an estimated genome size of 2 903 976 bp, knowing that 1 pg = 9.78×10^8^ bp (Doležel et al. 2003). A total of 268 plant samples were collected in Corsica, France, based on symptoms compatible with those caused by *X. fastidiosa;* 106 samples were collected in June 2017 and 162 in September 2018. For each French sample, DNA was extracted as described in PM7/24 (EPPO 2019) using two methods in order to optimize the chances of detection. CTAB-based extraction and robotic QuickpickTM SML kit from Bio-Nobile were used with the following modification: a sonication step (1 min, 42 KHz) was added after the samples (petioles, twigs) were finely cut, and was followed by a 15-min incubation period at room temperature. For initial laboratory diagnosis MLST results were compared with the Harper’s qPCR test (Harper et al. 2010) as in EPPO (2019) with following modifications: primers Xf-F and Xf-R, and probe Xf-P (Harper et al. 2010) were used at a final concentration of 0.6μM and 0.2μM respectively, non-acetylated BSA was used at final concentration of 1.5μg.μL^−1^, and 2 μl of DNA were used in 10μl reaction volume. Each DNA sample were tested in triplicates. To validate the nested PCR, DNA samples were provided by the National Reference Laboratory for Phytopathogenic Bacteria, Valencia, Spain, and from the Official Phytosanitary Laboratory of the Balearic Islands for determining *Xf* subspecies. Those DNA samples correspond to DNA extractions made from symptomatic plants sampled during official monitoring surveys. A total of 70 *Xf*-infected samples were analyzed from Balearic Islands and mainland Spain during 2018 (40 samples) and 2019 (30 samples), as well as 26 insect samples from both regions. DNA was extracted from petioles of symptomatic leaves as described in PM7/24 (EPPO 2019) using a CTAB-based extraction method for plant samples from Alicante and insect samples from Alicante and Balearic Islands. A Mericon DNeasy Food kit from Qiagen was used for plant samples from Balearic Islands. All DNA extraction methods have been validated; validation data is available in the EPPO Database on Diagnostic Expertise (EPPO, 2019).

### Nested-MLST primers and reactions

The seven HKG sequences (*cysG, gltT, holC, leuA, malF, nuoL, petC*) were extracted from 39 *Xf* genome sequences (S1 Table) (Denancé et al. 2019) to design the nested primers. Alignments were performed with BioEdit sequence alignment editor. The primers designed by Yuan *et al* (2010) were destined to be used as inner primers (PCR2) (Table 2) in our nested assay. We checked their characteristics with Primer3 V4.1.0 software (http://primer3.ut.ee/). Because of high Tm differences between forward and reverse primers for some primer pairs (*gltT*) (S2 Table), or high hairpin Tm values (*holC* forward primer), some primers from Yuan et al. (2010) were redesigned nearly at the same positions to improve their efficiency. Moreover, as primer sequences were already near the locus sequence ends, we also had to relocate some of them to design nested primers inside the sequence alignments without loss of informative sites. Outer primers (PCR1) were designed with Primer3 V4.1.0 software (http://primer3.ut.ee/) in flanking regions targeted by the inner primers. Outer and inner primers were tested *in silico* using a primer search tool available in the galaxy toolbox of CIRM-CFBP (https://iris.angers.inra.fr/galaxypub-cfbp) on 194438 bacterial Whole Genome Shotgun (WGS) sequences available in the NCBI database (as on March, 2019) including 58 *Xylella* and 1292 *Xanthomonas*, and *in vitro* on target and non-target bacterial strains (Table 1).

**Table 2.**
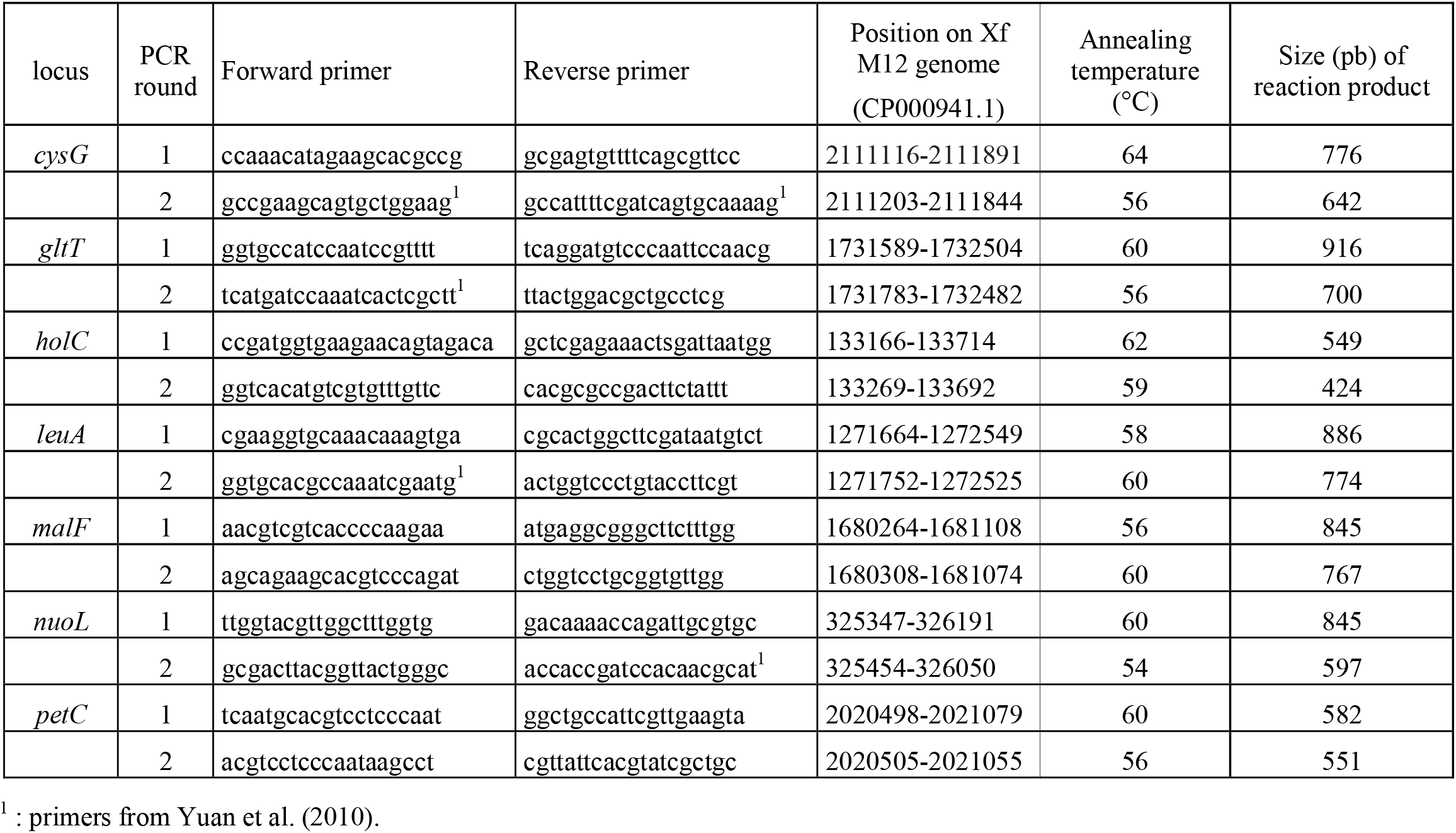
Primer sequences used in the *X. fastidiosa* nested-MLST scheme.

PCRs were performed in 25 μl reaction buffer (Promega) with MgCl2 at 1.5mM final, 200μM dNTP, 300μM each of the forward and reverse primers, 0.6 U GoTaq G2 (Promega) and 2μl of sample DNA. The first-round PCR program consisted of an initial denaturation step of 3 min at 95°C followed by 35 cycles of 30 s denaturation at 95°C, 30 s annealing at the relevant temperature according to each gene (determined by gradient PCR) and 60s elongation at 72°C followed by a final extension step of 10 min at 72°C (Table 2). The second round was performed with 30 cycles under same conditions and same concentrations but with a final volume of 50μl for sequencing purposes and with 4μl of first-round PCR product. The primer pairs of the second round of each nested PCR were used for sequencing (by Genoscreen, Lille, France for French samples and by Stabvida, Caparica, Portugal, for Spanish samples) the corresponding PCR products after 1.8% agarose gel visualization. To avoid contamination, one sample was opened at a time and stringent cleaning measures were applied after each experiment.

### Statistical analysis

The sensitivity of detection by conventional- and nested-MLST PCRs were compared in plant and vector samples for the seven HKGs that were analyzed by both approaches, by using a Chi square test using SAS (version 9.4, SAS Institute, Cary, NC, USA). Analysis was performed for the Spanish samples only, as HKG-PCRs were not systematically carried out on the French samples. Results were considered significantly different when *p* ≤ 0.05.

### Sequence acquisition, alignment and analyses

Forward and reverse nucleotide sequences were assembled, and aligned using Geneious 9.1.8 software (French samples) or Bionumerics V7.6.3 software (Spanish samples) to obtain high quality sequences. ST or loci assignation was performed according to http://pubmlst.org/xfastidiosa/. To reduce the costs of sequencing for French samples, only PCR products obtained for samples showing the highest rate of successful HKG amplifications were sequenced. On the other hand, all positive *holC* amplifications were sequenced to obtain a larger view of alleles present in Corsica.

## RESULTS

### Nested-MLST proved to be specific

The specificity of the outer and inner primer pairs was tested *in silico* and *in vitro. In silico*, all primers pairs showed the best scores of alignment with *Xf* genomic sequences. Some non-target organisms showed sequences nearly identical at outer primer locations with only one mismatch and a similar expected fragment size, but sequences of inner primers were more different indicating that there will be no amplification. This was the case for various *Xanthomonas* strains that contained one mismatch at position 15 of the *petC* forward outer primer and an identical sequence for the outer reverse primer. *X. taiwanensis holC* sequence corresponding to inner primers contained also only one mismatch. The fragment size predicted was as expected for *Xf*. Other predictions with one mismatch located in primers did not end in fragment amplifications of the same expected size. Then, the specificity of the outer and inner primer pairs (Table 2) was validated *in vitro* on five target strains and 33 non-target strains (Table 1). Specificity of the nested-MLST assay could not been tested *in vitro* on *X. taiwanensis* as no strain was available. Amplifications were obtained for all Xf strains. No amplification was detected on the non-target strains except for strain CFBP 2532 (*Xanthomonas oryzae* pv *oryzae*) and CFBP 2533 (*Xanthomonas hortorum* pv. *pelargonii*) in the first round of the nested PCR for the *petC* outer primers, providing a product of the expected size. However, these products were not amplified in the second round of the nested PCR and no false positive signal was finally obtained.

### Nested-MLST Limit of Detection is comparable to that of qPCR

The sensitivity of each primer combination was evaluated on serial dilutions of a genomic DNA solution calibrated (Qubit fluorimeter, Invitrogen) at 220 ng.mL^−1^ (Figure 1). First round PCRs gave a signal more or less intense for concentrations up to 2.2 ng.mL^−1^ (0.8 x 10^6^ copies.mL^−1^) for all HKG except *malF* and *cysG* (220 pg.mL^−1^). The second round of PCRs allowed a sufficiently strong signal for sequencing for concentrations up to 22 pg.mL^−1^ (0.8 x 10^4^ copies.mL^−1^) for *gltT, holC, petC, leuA, cysG*, and up to 2.2 pg.mL^−1^ (0.8 x 10^3^ copies.mL^−1^) for *nuoL* and *malF*. The same range of genomic DNA solutions was tested with the Harper’s qPCR test to compare sensitivity of these two tests (S3 Table). The latest signal (LoD) for the Harper’s qPCR test (Cq =37.64) was obtained with the concentration of 0.8.10^3^ copies.mL^−1^ and no amplification was obtained for lower concentrations.

**Figure 1 :**
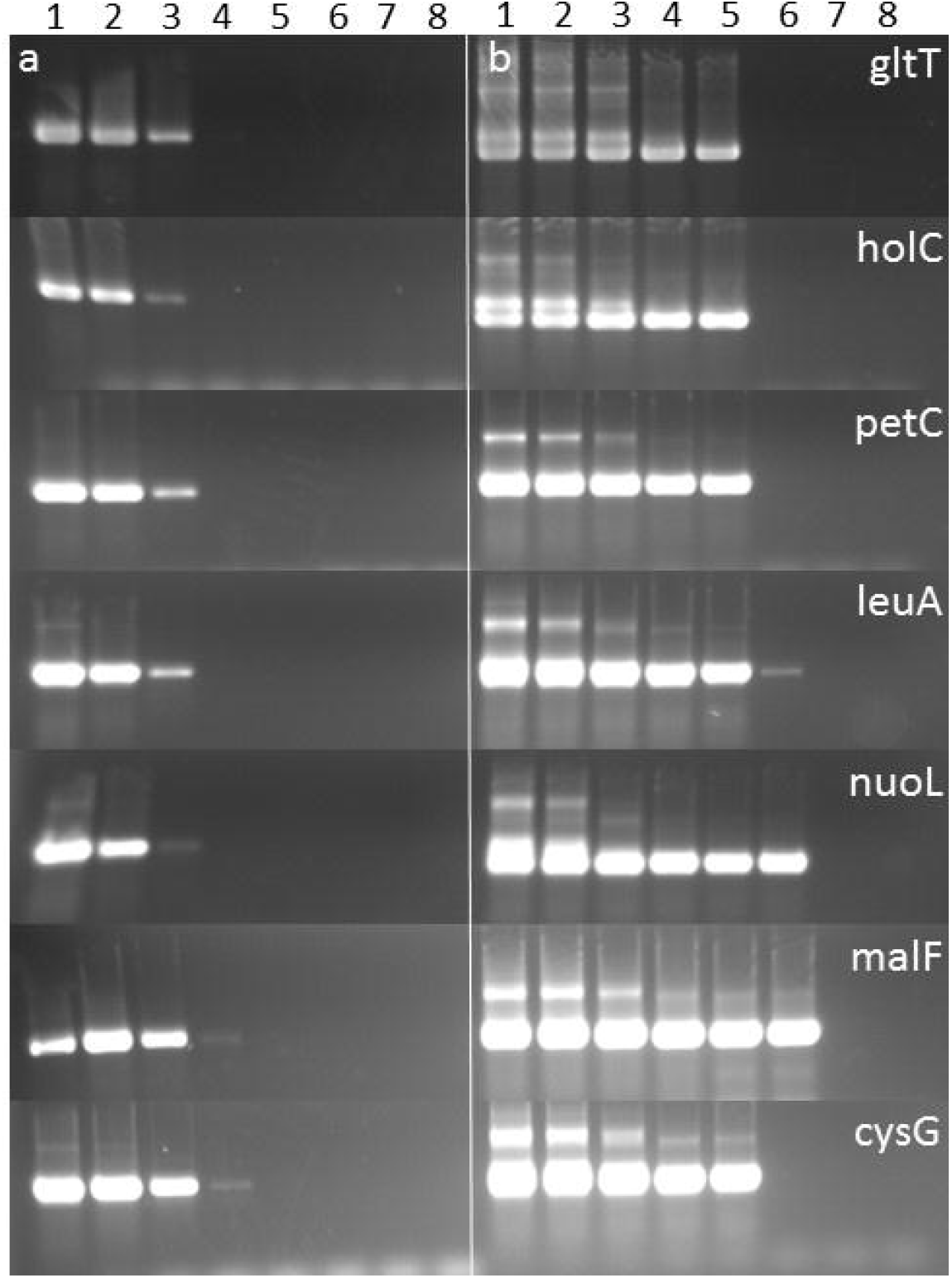
detection threshold of conventional-MLST (a) and nested-MLST (b) for seven HKGs using genomic DNA dilution range (1: 220 ng.mL^−1^; 2: 22 ng.mL^−1^; 3 : 2.2 ng.mL^−1^; 4: 220 pg.mL^−1^; 5: 22 pg.mL^−1^; 6: 2.2 pg.mL^−1^; 7 : 220 fg.mL^−1^; 8: 22 fg.mL^−1^)

Previously, we evaluated the LoD of the conventional PCRs for *cysG* and *malF* of the initial MLST scheme (Yuan et al, 2010) on a range of dilutions of CFBP 8070 genomic DNA with the Platinum Taq polymerase (Invitrogen) and tested the effect of adding BSA (final concentration at 0.3μg. μL^−1^) on the efficiency of the conventional PCRs. No improvement was obtained as all signals remained around 0.8 x 10^6^ bacteria.mL^−1^ (S1 Figure).

### Analysis of naturally infected samples

Using qPCR Harper’s test, 22 samples from 2017 and eight samples from 2018 collected in France were positive (Cq values<35) with one or both DNA extraction methods; 70 samples from 2017 and 36 samples from 2018 were equivocal (35≤Cq≤40), 14 samples from 2017 and 118 from 2018 were negative (Cq>40) (Table 3 and S4 Table). Positive and equivocal samples were tested using the first round of PCR of the MLST assay: five samples from 2017 (one *Spartium junceum*, three *Polygala myrtifolia*, and one *Genista corsica*) gave a signal for at least one gene, but no complete typing was obtained for any sample. No sample from 2018 gave a signal. Most of Spanish samples used to evaluate nested-MLST scheme were positive using Harper’s qPCR (only two out of 40 plant samples were equivocal in 2018 and eight out of 26 vector samples).

**Table 3:**
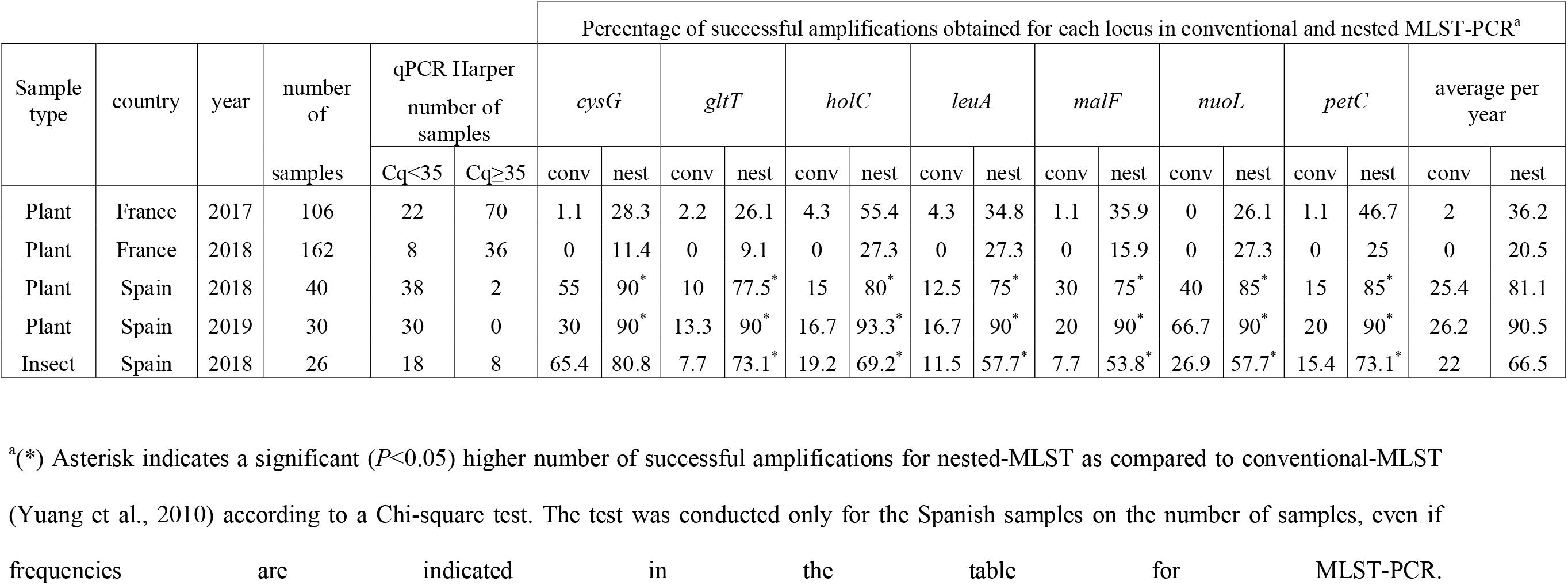
Number of samples, positive and equivocal in qPCR Harper. Percentage of successful amplifications obtained for each locus in conventional and nested PCR

### Nested-MLST improved successful HKG typing by increasing sensitivity level

Using nested-MLST for French samples, full allelic profiles were obtained for five samples from 2017 and one from 2018 corresponding to the lowest Cq in Harper’s qPCR test (Table 4 and S4 Table) Among fully typed samples, four were *X. fastidiosa* subsp. *multiplex* ST7 (*Genista corsica, Polygala myrtifolia, Spartium junceum*), and two were *X. fastidiosa* subsp. *multiplex* ST6 (*Polygala myrtifolia*).

**Table 4:** Allele numbers and STs obtained for fully typed samples in France and Spain for plant and insect samples. The numbers correspond to the names of the samples.

| Country | Sample names | *cysG* | *gltT* | *holC* | *leuA* | *malF* | *nuoL* | *petC* | sequence type (ST) |
| --- | --- | --- | --- | --- | --- | --- | --- | --- | --- |
| France | *Spartium junceum* 2 | 7 | 3 | 3 | 3 | 3 | 3 | 3 | ST7 |
| France | *Polygala myrtifolia* 3, 4 | 3 | 3 | 3 | 3 | 3 | 3 | 3 | ST6 |
| France | *Genista corsica* 1 | 7 | 3 | 3 | 3 | 3 | 3 | 3 | ST7 |
| France | *Polygala myrtifolia* 5, 6 | 7 | 3 | 3 | 3 | 3 | 3 | 3 | ST7 |
| Spain | *Cistus albidus* 2 | 31 | 15 | 10 | 7 | 17 | 16 | 6 | ST80 |
| Spain | *Ficus carica* 1 | 1 | 1 | 1 | 1 | 1 | 1 | 1 | ST1 |
| Spain | *Helichrysum italicum* 1 | 3 | 3 | 3 | 3 | 3 | 3 | 3 | ST6 |
| Spain | *Juglans regia* 1 | 1 | 1 | 1 | 1 | 1 | 1 | 1 | ST1 |
| Spain | *Lavandula angustifolia* 1 | 32 | 3 | 3 | 3 | 3 | 3 | 3 | ST81 |
| Spain | *Olea europaea* 1 | 3 | 3 | 3 | 3 | 3 | 3 | 3 | ST6 |
| Spain | *Phagnalon saxatile* 1 | 3 | 3 | 3 | 3 | 3 | 3 | 3 | ST6 |
| Spain | *Polygala myrtifolia* 1 | 3 | 3 | 3 | 3 | 3 | 3 | 3 | ST6 |
| Spain | *Prunus armeniaca* 1 | 3 | 3 | 3 | 3 | 3 | 3 | 3 | ST6 |
| Spain | *Prunus domestica* 1 | 32 | 3 | 3 | 3 | 3 | 3 | 3 | ST81 |
| Spain | *Prunus domestica* 2 | 3 | 3 | 3 | 3 | 3 | 3 | 3 | ST6 |
| Spain | *Prunus dulcis* 4-8,10,11,15,18-26,30-47 | 3 | 3 | 3 | 3 | 3 | 3 | 3 | ST6 |
| Spain | *Prunus dulcis* 9 | 31 | 15 | 10 | 7 | 17 | 16 | 6 | ST80 |
| Spain | *Prunus dulcis* 1,2 | 32 | 3 | 3 | 3 | 3 | 3 | 3 | ST81 |
| Spain | *Prunus dulcis* 3 | 7 | 3 | 3 | 3 | 3 | 3 | 3 | ST7 |
| Spain | *Rhamnus alaternus* 1 | 3 | 3 | 3 | 3 | 3 | 3 | 3 | ST6 |
| Spain | *Rosmarinus officinalis* 4 | 3 | 3 | 3 | 3 | 3 | 3 | 3 | ST6 |
| Spain | *Rosmarinus officinalis* 1,3 | 31 | 15 | 10 | 7 | 17 | 16 | 6 | ST80 |
| Spain | *Prunus domestica* 3 | 3 | 3 | 3 | 3 | 3 | 3 | 3 | ST6 |
| Spain | *Philaenus spumarius* 6,7,8,10,11 | 1 | 1 | 1 | 1 | 1 | 1 | 1 | ST1 |
| Spain | *Philaenus spumarius* 1 | 3 | 3 | 3 | 3 | 3 | 3 | 3 | ST6 |
| Spain | *Philaenus spumarius* 22 | 32 | 3 | 3 | 3 | 3 | 3 | 3 | ST81 |
| Spain | *Neophilaenus campestris* 1,2 | 3 | 3 | 3 | 3 | 3 | 3 | 3 | ST6 |

Our scheme was also evaluated on Spanish samples already proved infected by *Xf*. These samples from different outbreaks showed a wide range of Cq values ranging from 18.8 to 36.0 for plant samples and from 23.29 to 37.0 for insect samples (Table 3 and S4 Table). Samples were first analyzed using the conventional-MLST assay (Yuan et al. 2010). Amplification efficiency was variable and ranged from 10% for *gltT* to 67% for *nuoL* with an average of 25% and 26% for the seven HKG in 2018 and 2019, respectively. The nested-MLST assay improved the amplification efficiency that increased to 75% for *leuA* and up to 93% for *holC* with an average of 81% and 91% in 2018 and 2019, respectively. In total, full allelic profiles were obtained in seven plant samples using the conventional-MLST assay, whereas a total of 55 samples were fully typed with the improved nested-MLST assay (Table 4). For the 70 plant DNA samples that were tested by both protocols, for all the seven HKGs, conventional-MLST showed a significant (*P*<0.0005 for 2018 and *P*<0.0283 for 2019) lower number of samples amplified as compared to nested-MLST. Among fully typed plant samples using the nested-MLST, we identified *X. fastidiosa* subsp. *fastidiosa* ST1 in *Ficus carica* and *Juglans regia, X. fastidiosa* subsp. *multiplex* ST6 in *Helichrysum italicum, Olea europaea, Phagnalon saxatile, Polygala myrtifolia, Prunus armeniaca, Prunus domestica, Prunus dulcis, Rhamnus alaternus, and Rosmarinus officinalis*, *X. fastidiosa* subsp. *multiplex* ST7 in *Prunus dulcis, X. fastidiosa* subsp. *multiplex* ST81 in *Lavandula angustifolia and Prunus dulcis*, and *X. fastidiosa* subsp. *pauca* ST80 in *Cistus albidus, Prunus dulcis, and Rosmarinus officinalis*.

Not all insect samples could be tested by both protocols due to restrictions in DNA amount. In samples tested only by the original MLST assay (Yuan et al., 2010), the percentages of successful amplifications ranged from 8% (*gltT* and *malF*) to 65% (*cysG*). With the nested-MLST assay, successful amplifications ranged from 54% (*malF*) to 81% (*cysG*), with an average efficiency for the seven HKG of 22% to 67% for conventional and nested approach, respectively (Table 3, S4 Table). Nine insect samples were fully typed using a combination of both protocols (Table 4). *X. fastidiosa* subsp. *fastidiosa* ST1 was identified in insects from Mallorca (Balearic Islands), *X. fastidiosa* subsp. *multiplex* ST6 in insects from Alicante (mainland Spain) and *X. fastidiosa* subsp. *multiplex* ST81 in insects from Balearic Islands. For the nine insect samples that were tested by both protocols, conventional-MLST showed a significant (*P*<0.0247) lower number of samples amplified as compared to nested-MLST for six of the seven HKGs (excluding *cys*G). These results indicate that for insect samples it is also better to use directly the improved nested-MLST assay.

No nonspecific amplicons were observed in any of the samples. Negative controls (water) were run in the first and the second PCR and were always negative. The negative control coming from the first reaction always tested negative in the second one, confirming the absence of contamination during the entire process.. Positive control was a suspension of strain CFBP 8084 (ST29) from the subspecies *morus* or strain CO33 (ST72) as this STs were not previously found in Corsica, France or Spain, respetively.

### Nested-MLST allowed identification of new alleles among French samples

Incomplete profiles were obtained for various French samples due to variable amplification efficiencies varying according to the HKG. From 9% (with *gltT*) to 55% (with *holC*) of French samples gave a signal applying the nested-MLST assay. Alleles that were not yet described in plant samples in France were detected in 2017. This was the case for *holC*_1 and *holC*_2 alleles known to occur in ST from ST1 to ST5 and ST75 that cluster in the subspecies *fastidiosa* (https://pubmlst.org/xfastidiosa/). These alleles were sequenced in samples of *Asparagus acutifolius, Eleagnus, Cistus monspeliensis* and *C. creticus, Quercus ilex, Myrtus myrtifolia*, *Olea europea, Platanus, Arbutus unedo* (S4 Table). Other *holC* alleles already described in STs clustering in the subspecies *fastidiosa* (*holC*_24) were also sequenced from *Cistus monspeliensis and Pistaccia lentiscus. HolC*_10 alleles described in STs clustering in the subspecies *pauca* were sequenced from *Cistus monspeliensis* and *C. salicifolius, Cypressus, Metrosideros excelsa, Myrtus communis, Pistaccia lentiscus, Quercus ilex, Rubia peregrina, Smilax aspera* samples. Similarly, *holC*_3 (known in ST6, ST7, ST25, ST34, ST35, ST79, ST81 and ST87 clustering in the subspecies *multiplex*) were obtained from samples of *Acer monspeliensis, Arbutus unedo, Calicotome spinosa, Cistus monspeliensis, Genista corsica, Myrtus communis, Olea europea, Phyllirea angustifolia, Polygala myrtifolia, Quercus ilex and Q. pubescens, Spartium junceum*. Among samples from 2018, only *holC*_1 allele was detected in *Olea europea, Quercus ilex*, and *Platanus sp*. samples, and *holC*_3 allele in *Cistus monspeliensis*, *Acer monspeliensis, Myrtus communis, and Polygala myrtifolia* samples.

### Recombinants or mixed infections were identified by nested-MLST

Some French samples were further sequenced for several loci and these sequencing confirmed the presence of alleles occurring in the subspecies *fastidiosa, multiplex* and *pauca* (S4 Table). All alleles were previously described but were detected in combinations that were not previously described, suggesting the presence of recombinants or of mix infections (S4 Table). This is the case for *Cistus monspeliensis* 7 showing an unknown combination of *cysG*_2/ *petC*_2/ *nuoL*_2/ *gltT*_2 (known in ST5) with *malF*_4 (known in ST2), both from subspecies *fastidiosa*; *Helichrysum italicum* 1 showing *leuA*_1 (known in subspecies *fastidiosa*) with *petC*_3/ *holC*_3 known in subspecies *multiplex; Myrtus communis* 4 with *leuA*_3/*holC*_2 respectively known in subspecies *multiplex* and *fastidiosa*; *Myrtus communis* 8 and *Platanus* presenting form 1 alleles for five HKG mixed with *malF*_4 (all known in subspecies *fastidiosa*) and *Q. ilex* 10 presenting form 1 alleles for two HKG mixed with *malF*4); *Olea europaea* 2 with four *multiplex* alleles combined with *nuoL*_1 (subspecies *fastidiosa*); *Olea europaea* 5 with four *pauca* alleles combined with *malF*_15 (known in ST72 and ST76, subspecies *fastidiosa*). Two samples gave a double sequence for *holC* that were impossible to analyze (S4 Table). Some sequences were ambiguous with superimposed peaks at some locations in otherwise good quality chromatograms revealing mixed infections. In those 12 samples, the number of potential combinations was too high to detect one probable allelic form, excepted for *Prunus dulcis* where the superimposed chromatograms corresponded to only two allelic forms (*holC*_3 or *holC*_6 which are found in subspecies *multiplex*). The *holC*_6 allelic form and the *leuA*_5 allele obtained for this sample are found in ST10, ST26, ST36, ST46, and ST63.

## DISCUSSION

A two-step nested procedure for MLST was developed to improve the typing of samples infected with low *Xf* population sizes that cannot be typed using the conventional protocol. In order not to affect the comparability of the results with the databases, the widely used MLST scheme developed for *Xf* that is supported by the pubMLST public website (Yuan et al. 2010) was re-used.

The nested-MLST approach proved to be specific and efficient. No nonspecific amplifications were observed in any of the samples. Moreover, the sensitivities of the Harper’s qPCR detection test and the nested-MLST were similar with a LoD ranging from 10^3^ bacteria.mL^−1^ to 10^4^ bacteria.mL^−1^ These LoDs are similar to other nested-MLST approaches such as those developed for *Burkholderia cepacia* (Drevinek et al., 2010) but higher than for the one developed for *Neisseria meningitidis* (10 copies mL^−1^) (Diggle et al, 2003). Consequently, in resource-limited settings where qPCR facilities are not available, the assay may be used as a useful diagnostic tool if applied with all necessary precautions to avoid cross-contamination between samples. The sequencing, which is costly, can be done as a consecutive but separate step to provide information on subspecies present in the sample. Higher bacterial loads (as indicated by lower Cq values) were observed in Spanish samples than in French samples, for which low amplification efficiency and partial profiles were observed. Full allelic profiles (ST6 and ST7 from *multiplex* subspecies) were obtained for *Polygala myrtifolia*, *Spartium junceum* and *Genista corsica* samples from France probably because they carried a higher bacterial load as shown by the low Cq obtained with the Harper’s qPCR test: five of the six typed samples had a Cq value between 23.4 and 26.5. The use of the nested-MLST assay to type plant Spanish samples allowed a higher number of successful complete typing (55 samples versus seven samples with the conventional approach). Spanish samples generally showed higher *Xf* titer (i.e, lower Cq values in Harper’s qPCR test) than the French samples but also concerned different plant species.

In our nested-MLST assay as well as in the original MLST assay, the amplification efficiencies were variable among genes, while all primers were designed using the same parameters from the software. For example, the *holC* gene for French samples tested with the nested-MLST assay was successfully amplified in 55% of samples collected in 2017 while the *gltT* and *nuoL* genes gave the lowest rates (around 26%). For samples collected in Spain tested with the original MLST assay, amplification rates among the seven HKGs ranged from10 to 67%. Success rate variations were also observed in medical research using MLST between samples and between loci (Weiss et al. 2016). When conducted on strains, no differences about amplification rates are observed because of DNA excess. Robustness of a PCR reaction is determined by appropriate primers and it is not always obvious why some primer combinations do not amplify well, even if some parameters such as DNA folding can interfere in PCR efficiency (Bustin & Huggett 2017). In this study, even if primer annealing temperature was adjusted, design of primers was limited by their arbitrary localization.

Typing results of French samples were concordant with previously published results (Denancé et al. 2017) but also revealed the presence of alleles not yet described in France. It should be noticed that no unknown sequence was obtained, refraining from evoking contaminations as the origin of these yet undescribed alleles in France. Thanks to the high rate of amplification of *holC* in nested PCR, it was also possible to obtain sequences for equivocal samples (Cq with the Harper’s qPCR test above 35) to confirm the presence of the bacterium in these samples. Surprisingly, these amplifications led to alleles that correspond to subspecies other than the *multiplex* subspecies. Thereby, alleles from subspecies *pauca* (*holC*_10) and *fastidiosa* (*holC*_1, *holC*_2, *holC*_24) were sequenced. *HolC*_10 was already reported in *Polygala myrtifolia* in the south of France in 2015 (Denancé et al., 2017). *HolC*_1 finding is in agreement with Cruaud et al (2018), who also reported *holC*_1 in insects in Corsica. Up to now, no *holC*_2 was reported in France but it is known in the USA. *HolC*_24 was also reported in *Polygala myrtifolia* in Corsica in 2015 (Denancé et al. 2017). Further plant sampling efforts are needed to confirm the establishment of those strains in the environment or to document further the dynamics of alleles revealing sporadic infections.

For French samples only, several samples could not be typed since the chromatograms showed an overlap of two peaks precisely on the polymorphic sites (mainly with *leuA* and *holC* genes). This has already been reported by Denancé et al (2017), it suggests the simultaneous presence of several strains in the same sample since only one copy of these genes are known in *Xf* (Yuan et al, 2010). Moreover, the report of previously unknown combination of alleles belonging to different subspecies can also results from the presence of co-infection or of recombinants. Recombination events are reported in *Xf* (Denancé et al. 2017; Jacques et al. 2016, Nunney et al. 2014a, Saponari et al. 2019) and could have led to host shift (Nunney et al. 2014b). In this study, eight samples presented unknown combinations of alleles from the same or different subspecies which could be explained by intrasubspecies or intersubspecies recombination events. As reported in Potnis et al. (2019), such events may exist and occur but not with the same frequency. Moreover, natural competence can be variable among *Xf* strains (Kandel et al., 2017). These events could also reflect a mechanism of adaptation (Kandel et al., 2016). Five samples among these eight samples were collected in 2017 and three in 2018, and were different between years. In 2018 the three cases were a similar combination of alleles and were found in three different plants. Future surveys will be necessary to know if some of these recombinants strains are indeed present in Corsica or are the consequence of mixed infections and if they have adapted and survived on different hosts.

The objective of this study was to improve the published MLST scheme supported by a public website (https://pubmlst.org/xfastidiosa/) by designing nested primers to lower the limit of detection and help in *Xf* diagnosis and typing. Thus, this improved MLST assay enables a higher sensitivity and specific typing of *Xf* directly from plant and insects samples without the need of isolating the strain and at an affordable cost.

## Author Contributions

SC, QB and MMB, MPVA performed the experiments, ED took part to primer design, MB helped with bioinformatic tools, SC conceived the study, MAJ and BL applied for funding, SC, ED, MAJ, BL wrote the manuscript. All authors read and approved the final version of the manuscript.

## Acknowledgments

We thank Muriel Bahut (ANAN technical facility, SFR QUASAV, Angers, FR) for DNA extraction automatization, CIRM-CFBP (Beaucouzé, INRA, France; http://www6.inra.fr/cirm_eng/CFBP-Plant Associated-Bacteria) for strain preservation and supply. We thank Ester Marco-Noales from the National Reference Laboratory for Phytopathogenic Bacteria (IVIA), and Diego Olmo from the Official Phytosanitary Laboratory of the Balearic Islands for providing DNA samples for MLST typing.

## Nomenclature

BLAST: Basic Local Alignment Search Tool
Cq: quantification cycle
HKG: housekeeping gene
INRA: French National Institute for Agricultural Research
IRHS: Research Institute of Horticulture and Seeds
LoD: Limit of Detection
MLST: Multilocus Sequence Typing
NCBI: National Center for Biotechnology Information
ST: Sequence Type
*Xf*: *Xylella fastidiosa*
WGS: Whole Genome Shotgun

## Supplementary Material

S1 Table: List of *X. fastidiosa* genome sequences used in this study for primer and probe design (Denancé et al. 2019)

S2 Table : Primers properties

S3 Table: detection threshold for Harper’s qPCR test using genomic DNA dilution range (1: 220 ng.mL^−1^; 2: 22 ng.mL^−1^; 3 : 2.2 ng.mL^−1^; 4: 220 pg.mL^−1^; 5: 22 pg.mL^−1^; 6: 2.2 pg.mL^−1^; 7 : 220 fg.mL^−1^; 8: 22 fg.mL^−1^)

S4 Table: results obtained with qPCR and nested-MLST. (+) means that a signal has been obtained in PCR but the PCR product has not been sequenced.

S1 Figure: detection threshold of conventional PCR for *cysG* and *malF* loci (Yuan et al., 2010) with and without BSA (final concentration at 0.3 μg. μL^−1^) using genomic DNA dilution range (1: 220 ng.mL^−1^; 2: 22 ng.mL^−1^; 3 : 2.2 ng.mL^−1^; 4: 220 pg.mL^−1^; 5: 22 pg.mL^−1^; 6: 2.2 pg.mL^−1^; 7 : 220 fg.mL^−1^; 8: 22 fg.mL^−1^). (+) positive control; (-) negative control.

