## Supplementary figures and images for "Development of a Nested-MultiLocus Sequence Typing approach for a highly sensitive and specific identification of *Xylella fastidiosa* subspecies directly from plant samples"

### supplemental Figure 1

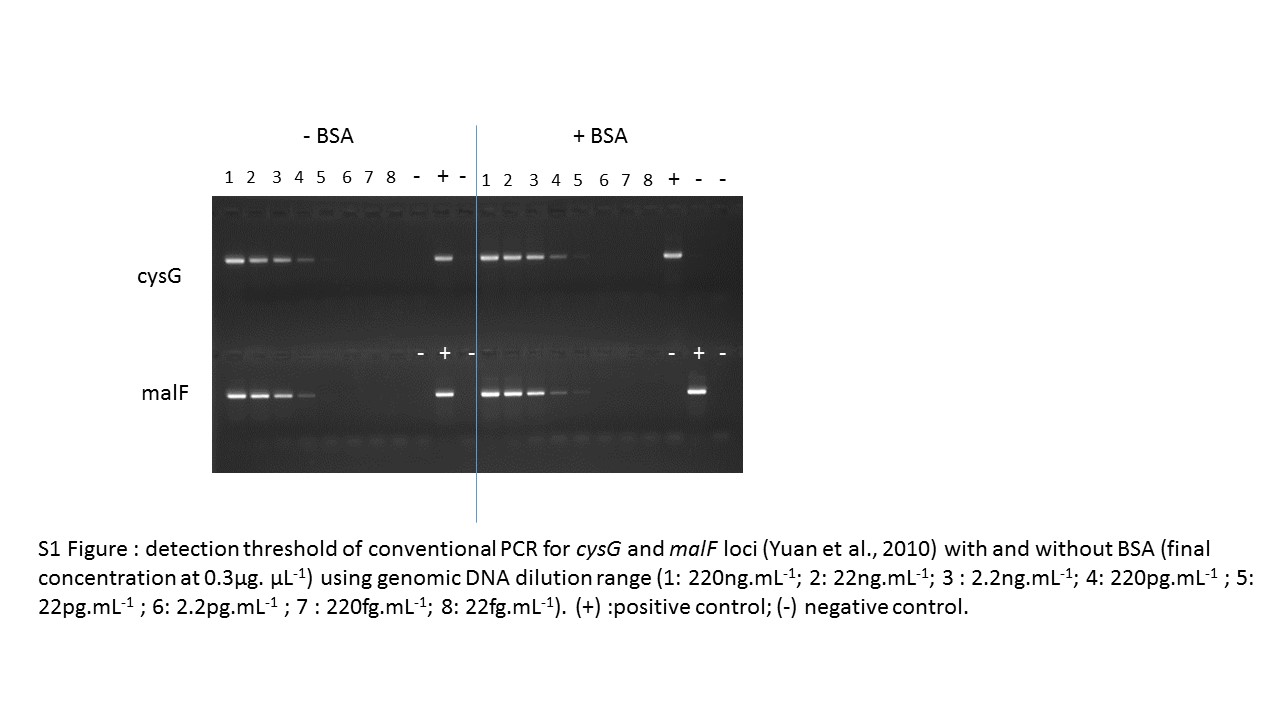
